# HiCArch: A Deep Learning-based Hi-C Data Predictor

**DOI:** 10.1101/2021.11.26.470146

**Authors:** Xiao Zheng, Jinghua Wang, Chaochen Wang

## Abstract

Hi-C sequencing analysis is one of the most popular methods to study three-dimensional (3D) genome structures, which affect the gene expression and other cellular activities by allowing distal regulations in spatial proximity. Hi-C sequencing analysis enhances understanding of chromatin functionality. However, due to the high cost of Hi-C sequencing, the publicly available Hi-C data of high resolutions (such as 10kb) are limited in only a few cell types. In this paper we present HiCArch, a light-weight deep neural network that predicts Hi-C contact matrices from 11 common 1D epigenomic features. HiCArch identifies topological associated domains (TADs) of 10kb resolution within the distance of 10Mb. HiCArch obtains train Pearson correlation score at 0.9123 and test Pearson correlation score at 0.9195 when trained on K562 cell line. which are significantly higher than previous approaches, such as HiC-Reg[1], Akita[2], DeepC[3], and Epiphany[4].

## 1 Introduction

Three-dimensional (3D) genomic structure has been considered as a critical genomic feature over the past two decades[5, 6, 7, 8], because the spatial distance between the regulatory elements and the genes affects the probability they interact with each other, which further influence gene expression. Abnormal 3D genomic structure can mediate genetic variants associated with long-range regulatory interactions, leading to common diseases such as Amyotrophic lateral sclerosis and Crohn’s disease[9, 10]. To study 3D genomics, researchers have developed various technologies which involve microscopic imaging technology such as FISH[11], and chromosomal conformation capture technology including 3C[12], 4C[13, 14], 5C[15], CHIA-PET[16], and Hi-C[17]. Among these technologies, Hi-C is a next-generation sequencing based approach that provides the frequencies of all chromosomal contacts at once and requires fewer experimental skills. However, owing to high sequencing cost, Hi-C data of high resolutions (such as 10kb) are far less obtainable than other epigenomic data, such as those generated from the ChIP-seq assays.

Previous studies have utilized machine learning methods to predict Hi-C contact matrices with sequencial data. For example, DeepTACT[18], DeepC[3] and Akita[2] extract features from DNA sequences and generate the Hi-C maps with convolutional neural networks, and HiCReg[1] uses Random Forests regression to predict contact frequencies pixel by pixel. However, DeepTACT, DeepC and Akita lack generalization capability to different cell types, and are dependent on massive DNA inputs; in contrast, HiCReg learns the features with epigenetic marks only but misses localized pattern between the contact pairs, and it could only generate the interaction between one contact pair at once, reducing the predicting efficiency. Few methods could capture useful features from the whole epigenomic sequences only, followed by 2D Hi-C contact map generation and optimization.

In 2019, the natural language processing community developed transformer[19] a deep learning model architecture which represents the long range context by computing the interactions between each position. Its nature of describing pair-wise interactions within the input sequences, with the advantages of more parallelizable and requiring less training time than the traditional architectures like convolutional neural networks (CNNs) and recurrent neural networks (RNNs), transformer model has been applied to infer biomolecular structures based on sequences, such as AlphaFold2[20] for protein structure prediction from amino acids, and E2Efold[21] for RNA secondary structure prediction from transcriptional base pairs, which implies potentials of using similar deep learning models to predict Hi-C from epigenetic data.

Hence, we proposed HiCArch, a transformer-based model architecture for Hi-C contact matrices prediction based on the 11 types of K562 epigenomic features, consisting of chromatin binding factors and histone modifications. HiCArch processes the sequential input and generates the 2D Hi-C matrix via two main modules: sequence-to-sequence (seqToSeq, or STS) module, sequence-to-matrix (seqToMat, or STM) module. While the STS module extract sequencial patterns from inputs, the STM module expands the STS-generated sequences to contact matrices by pair-wise dot product. To test the performance of transformer model, we incorporated transformer layers as well as CNN layers into STS. By comparing STS with various proportion of transformer layers (0, 50%, 100%), we demonstrated hybridization 50% transformer layers following 50% CNN layers generated the model with highest train and test accuracies. With customized loss function design and optimization, our final HiCArch model can accurately predict chromatin contacts within the window of 10Mb, with the resolution of 10kb, equivalent to the input features. Also, HiCArch made acceptable prediction on another cell line GM12878, indicating its high generalization capability. Moreover, as no DNA sequences are required, it is convenient for other researchers to customize HiCArch on their interested cell types.

## 2 Methods

### 2.1 Dataset Settings

The Hi-C matrices of 10kb resolutions were downloaded from the GEO dataset with the accession number GSE63525[22]. For the 1D epigenomic data, the .bam files aligned to hg19 genome assembly were downloaded from ENCODE project (https://www.encodeproject.org/). The 11 epigenomic features consist of the chromatin binding factors (CTCF, TBP), chromatin accessibility (DNase I), and histone modifications (H3K4me1, H3K27ac for enhancer signatures, H3K36me3, H3K79me2, H4K20me1, H3K4me3 for gene activation, and H3K9me3, H3K27me3 for gene repression). For each feature, we used SAMtools[23, 24] to merge the highly correlated replicates. Based on these samples, tag directories were created by Homer makeTagDirectory[25]. The bedGraphs binned at 10kb resolution were then created from the tag directories by Homer makeUCSCfile. To balance the contribution of all features within the chromosome, we dropped records at the beginning and the end of each chromosome where one or more features were absent. To avoid the effect of extremely large outliers, we replaced them with smaller pseudo values that were still larger than normal. To fit the data for HiCArch, we reshaped the data into an *N* × *b* dataframe, with N rows to denote the total number of records and 12 columns to represent the starting positions of each bin (per 10kb) and the 11 epigenomic features. Finally, the records of different chromosomes were stored in separate files (e.g. chr1_10kb_epi.txt in the sample data).

Our dataset consist of randomly selected sequences of epigenomic data with length 1000 and corresponding Hi-C matrices with size 1000 × 1000. Each sample is generated from one chromosome without data padding, and different samples can overlap in their data segments. The probability of a sample being drown from a certain chromosome is proportional to its DNA sequence length, and every sub-sequence has the same probability of being selected as a sample within a chromosome. The dataset sample input is an epigenomic sequence with shape (11, 1000), and the data sample output is a dense matrix with shape (1000, 1000). In practice, we provide dataset samples in batches with variable size from 8 to 16. All the data samples are normalized per-chromosome: the data sample inputs are normalized feature-wise into the range of [0, 1], while the data sample outputs are normalized into [0, 1] by dividing the original Hi-C data by the largest contact values in the Hi-C matrix of each chromosome. Considering the availability of both epigenomic and Hi-C data, the commonly used cell lines K562 and GM12878 were selected as HiCArch inputs in this work. We set chromosome 1-17 of K562 as training dataset, and chromosome 18-22 and chromosome X of K562 and GM12878 as test dataset.

Besides the above normalization steps, we added an extra Hi-C matrix per-distance-to-diagonal normalization to address a Hi-C matrix data distributive pattern of data values respect to the distances to the main diagonal. For a Hi-C matrix *M* ∈ ℝ^*l×l*^, the value of *M*_*j,k*_ quickly decreases as the distance to main diagonal |*j* − *k*| increases, making the values near the main diagonal being significantly larger than the values far off main diagonal. This pattern makes it difficult for deep learning models to extract numerical relationship between epigenetic sequences and Hi-C matrices. To alleviate this effect, we proposed a position-wise weight function 𝒲 : ℝ^*l×l*^ → ℝ^*l×l*^ which is defined as:

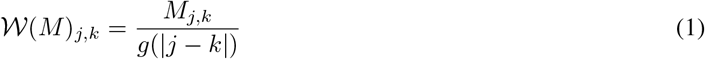

where 1 ≤ *j* ≤ *l*, 1 ≤ *k* ≤ *l, M* ∈ ℝ^*l×l*^, and *g* : ℝ → ℝ is our fitted Gradient Boost Regression model for per-distance-to-diagonal normalization on Hi-C data, which is fitted on multiple chromosomes in the K562 cell line.

### 2.2 Model Architecture

Our HiCArch model can be partitioned into two components: sequence-to-sequence processing component (seqToSeq or STS), and sequence-to-matrix processing component (seqToMat or STM). STS component accepts input epigenetic data with 11 features of length *l*, and generates a processed sequence with *d* pseudo-features of the same length *l*; STM component accepts the processed sequence with *d* -features from STS, and transforms it to an *l* × *l* output matrix. From the biological perspective, epigenetic data are “vague” features which do not contain the exact DNA base pair sequences, and Hi-C data are also “vague” due to the absence of detailed three dimensional structure. Unlike protein folding, the prediction from epigenetic data to Hi-C data is closer to a more generalized sequence-to-matrix prediction with much fewer geometric and spatial constraints. From the perspective of data dimensions, theoretically we separate our model into three phases: sequence-to-sequence, sequence-to-matrix, and matrix-to-matrix. Due to the computational complexity and memory cost restrictions, we didn’t use matrix-to-matrix computation component in practice.

The STS component is composed of multiple blocks of 1-d convolution layers and transformer layers[19] for 1-d sequential data processing. We have experimented with different architectures using multiple settings including stacking only 1-d convolution layers, stacking only transformer layers, and stacking 1-d convolution layers followed by transformer layers. We have also included multiple experiments with layer-wise residual connections or block-wise residual connections similar to ResNet[26]. For simplicity we keep our STS component simple and straightforward with sequential architectures and relatively shallow depth (within 20 layers). When transformer layers are used, the positional embedding is added at the first transformer layer.

The STM component is composed of our customized 1-d to 2-d convolution operator (upDimConv), a batch normalization layer[27], and a ReLU activation layer. Our UpDimConv operator is defined as:

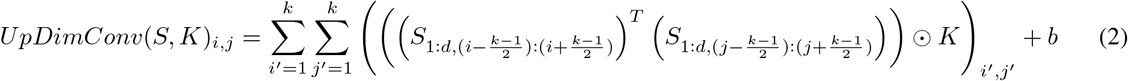

where *S* ∈ ℝ^*d×l*^ is the input to the operator, *K* ∈ ℝ^*k×k*^ is the symmetric kernel weight parameters such that *K* = *K*^*T*^, *b* ∈ ℝ is the bias parameter, *k* is the kernel size, which is a positive odd integer.

The UpDimConv operator includes localized features in the input sequence and converts them to output matrix with appropriate data dependence patterns. 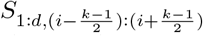 represents a zero-padded 1-d sliding window centered at *i*. The matrix multiplication of the two sliding windows captures the localized features in the form of linear combinations, and the localized features are further weighted by the symmetric kernel weight parameters. With the final bias addition step, this operator achieves a light-weight feature mapping from 1-d sequence to 2-d matrix with localized contact patterns which we expect our final model can precisely capture using trained parameters.

### 2.3 Accuracy Metrics

Although plenty of loss function formulations for structural matrix predictions are already available, they are mostly designed for solving three dimensional structure problems with direct spatial constraints like protein folding[20] or RNA secondary structures[21], which do not fit into our problem. With no direct spatial constraint obtained from Hi-C matrices, our case is similar to the plain symmetric matrix comparison problem that is different from comparisons of structural matrices with spatial implications. In the same research domain with aims of predicting Hi-C data from epigenetic data, HiCReg[1] uses a sequence-to-point strategy, which is different from our sequence-to-matrix strategy; epiphany[4] proposed adversarial loss in their approach, but the adversarial loss configuration decreases its model inference accuracy in Pearson Correlation Coefficient by providing sharper but more inconsistent predictions with the ground truth, when compared with their mean square error loss configuration. Instead of using existing loss formulations, we proposed our own loss function ℒ : (ℝ^*l×l*^, *ℝ*^*l×l*^) → ℝ defined as:

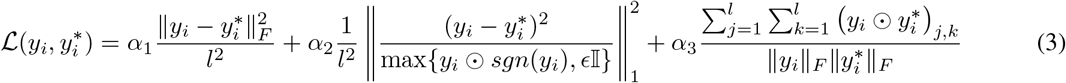

where *y*_*i*_ ∈ ℝ^*l×l*^ is the *i*-th sample prediction, 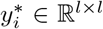 is the *i*-th ground truth output, *l* is the sequence length of each sample, which is 1000; 𝕀 is the *l* × *l* all one matrix, *E >* 0 is a small scalar for avoiding division by zero, *sgn* is the *l* × *l* dimension sign function; ‖ · ‖_*F*_ is the Frobenious norm, ‖ · ‖_1_ is the *L*_1_-norm, ⊙ represents the element-wise multiplication; *α, β, γ* ∈ [0, 1], and 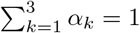.

As stated above, our loss function is composed of three terms: mean square error (MSE) term with weight *α*_1_, position-wise weighted term with weight *α*_2_, and cosine similarity term with weight *α*_3_. Note that the cosine similarity term can also be written as 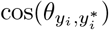 where 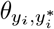 is the angle between *y*_*i*_ and 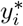 in the *l* × *l*-dimension Euclidean space. MSE term guarantees that the means and standard deviations of *y*_*i*_ and 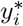 are close; position-wise weighted square term reduces the position-wise “outlier” predictions, especially the false large values on the locations far from the diagonal in our Hi-C matrix prediction where no large contacts should be predicted; and the cosine similarity term guides the model to provide similar distribution patterns to the ground truth in its model inference results.

When measuring the model inference accuracy, Pearson Correlation Coefficient is used. To be precise, given prediction *y* ∈ ℝ^*l×l*^ and ground truth *y*^*∗*^ ∈ ℝ^*l×l*^, the Pearson Correlation Coefficient is:

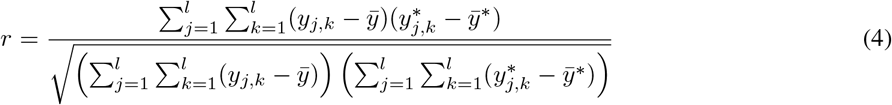

where 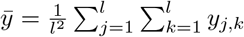, and 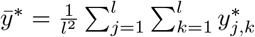.

Our accuracy formulation is almost the same as HiCReg[1] and epiphany[4], which guarantees fair comparisons among our HiCArch model and their solutions.

## 3 Results

### 3.1 The Deep Learning Approach Overview

Although many studies have been done to produce 2D matrices from 1D sequences, most of them either require exact sequences for precise predictions, or depend on structural constraints/reconstruction, such as protein folding and RNA secondary structure prediction. Unlike previous work, HiCArch aims to predict Hi-C matrices using only the epigenomic data with the same resolution as outputs, which is a vague-to-vague prediction. Also, little prior knowledge is available about structural constraints of Hi-C contacts. Therefore, we have to design a novel deep learning model for this task instead of just borrowing existing solutions.

Figure 1 is an overview of HiCArch. The input data of length *l* with *c* channels firstly undergo sequence processing by the STS module. STS consists of convolutional and/or transformer layers as needed. If both layer types are used, the convolutional layers are stacked over the transformer layers to ensure the output data shape. Then, the data of shape (*d, l*) are expanded into matrices (*l, l*) using the STM module. For each pair of positions across *l* × *l*, STM extracts their local features by convolution, which are further multiplied together to generate the predicted contact between these two positions. To evaluate the model, we calculate the loss using a formulation with three customized components, as described in Method. Then, the gradient is calculated with the loss and reduced by Adam[28], which is a popular gradient decent function that efficiently refines the deep learning model with an adaptive learning rate.

**Figure 1:**
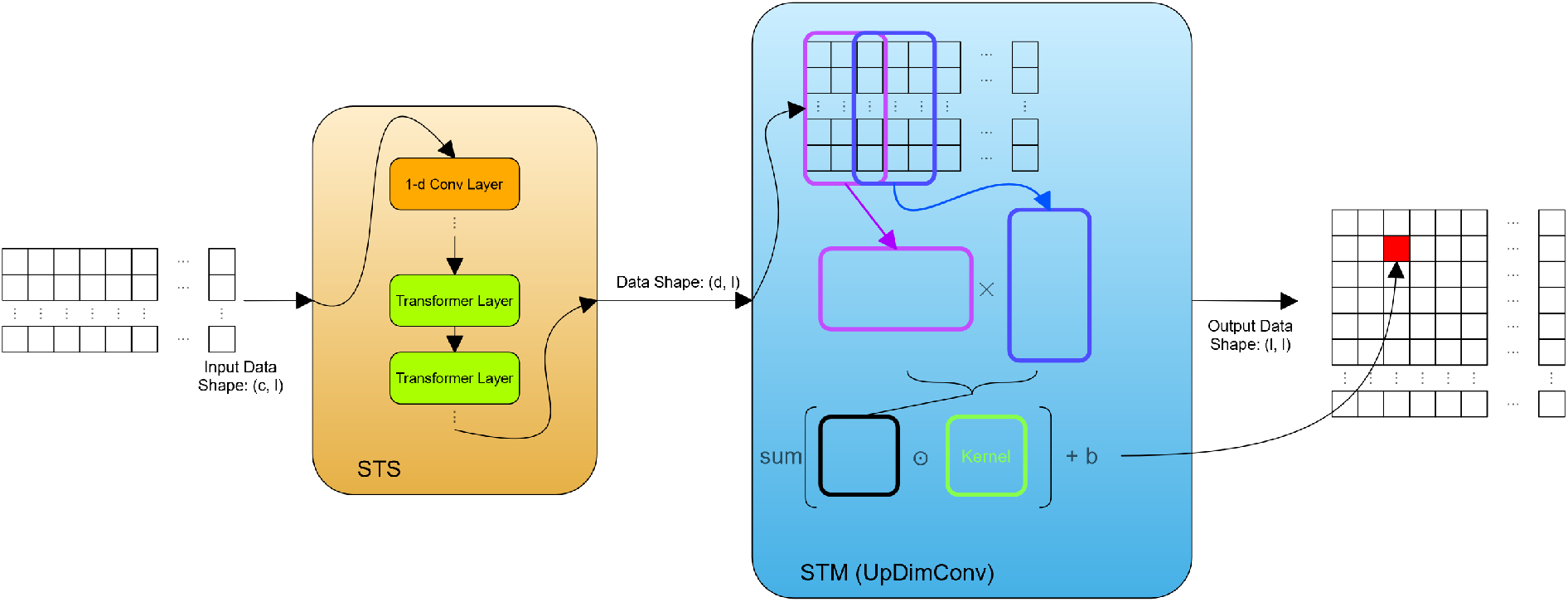
A brief data-flow of HiCArch

### 3.2 Model Settings

To determine whether transformer would give better prediction than CNN, we incorporated STS with 8 CNN layers or STS with 8 transformer layers into HiCArch and compared their performance. Unfortunately, due to the frequent contacts between nearby sites, all the predicted matrices only had values near the diagonal while missed distal interactions. We found that the mean contacts followed similar pattern with distance to diagonal across chromosomes (Figure 2a), so a curve averaging each chromosome was fitted using Gradient Boost Regression, which was applied to the data for balancing the distance-dependent effect (Method).

**Figure 2:**
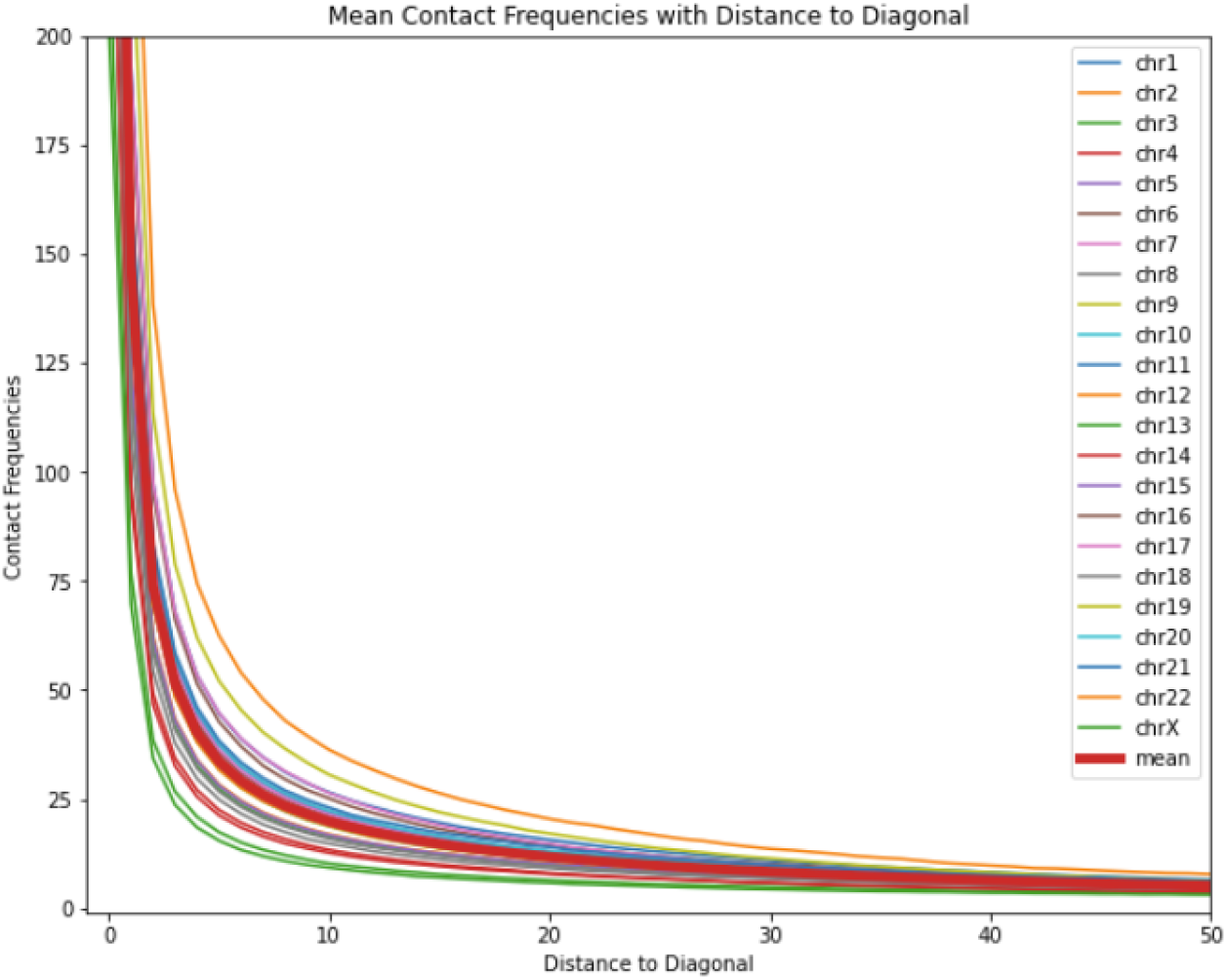
Usage of per-distance-to-diagonal normalization was validated by the similar contact patterns observed across chromosomes of K562.

As a result, STS (with pure convolutional layers) + STM has higher Pearsonr than STS (with pure transformer layers) + STM (Table 1), but the former produced more ambiguous matrices than the later (Figure 2b). To combine the advantages of CNN and transformer, a third STS with 50% convolutional layers and 50% transformer layers was designed. As expected, this configuration has a higher prediction accuracy than with transformer layers only or convolution layers only (Table 1), and its predicted matrices were visually more similar to the true matrices than the other two (Figure 2b). One possible explanation is that the convolution layers could extract local features, which were further handled by the transformer layers by highlighting the global structures. Given its superior performance than the other two models, this STS (with 50% convolutional layers and 50% transformer layers) +STM structure was used in the following experiments.

**Table 1:**
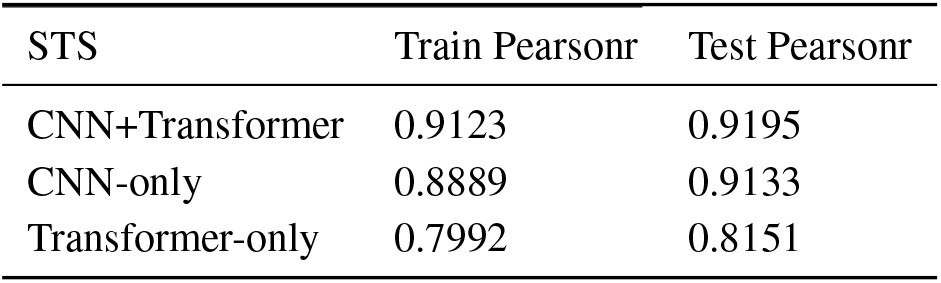
Model Accuracy Comparison on different STS

| STS | Train Pearsonr | Test Pearsonr |
| --- | --- | --- |
| CNN+Transformer | 0.9123 | 0.9195 |
| CNN-only | 0.8889 | 0.9133 |
| Transformer-only | 0.7992 | 0.8151 |

### 3.3 Loss Settings

In the Methods section, we defined the loss formulation as a combination of three terms: MSE, position-wise weighted square, and cosine similarity, with the corresponding weights to be *α*_1_, *α*_2_, *α*_3_ respectively. The weights of each term should be controlled finely, otherwise the predicted matrices would contrast sharply to the truth. For example, if overlarge cosine similarity is used, the resulting matrices might be visually similar to truth but with incorrect sign symbols; however, if the position-wise weighted square term is too large, blank matrices can be resulted, because most positions in truth are zeros. Therefore, using the STS (CNN & transformer) + STM model described previously, we calibrated the weights in the loss formulation so that the predicted matrices were not only looked like the true Hi-C matrices, but also similar in overall range as well as values at each position. The following results only presented models with slight tunning of each weight, as further changes largely reduced the model performance.

Compared with the basic settings previously used for STS selection, it was shown that loss settings with *α*_1_ = 0.09, *α*_2_ = 0.01, *α*_3_ = 0.9 improved the test accuracy by 0.06%. Also, the model under this loss setting could better distinguish strong or weak interactions, and predicted less false positives (Figure 3). Importantly, its predicted matrices clearly showed topological domains (blue blocks in Figure 3), and distal interactions (blue dots pointed by red arrows in Figure 3), indicating reliable biological discoveries using our model.

**Figure 3:**
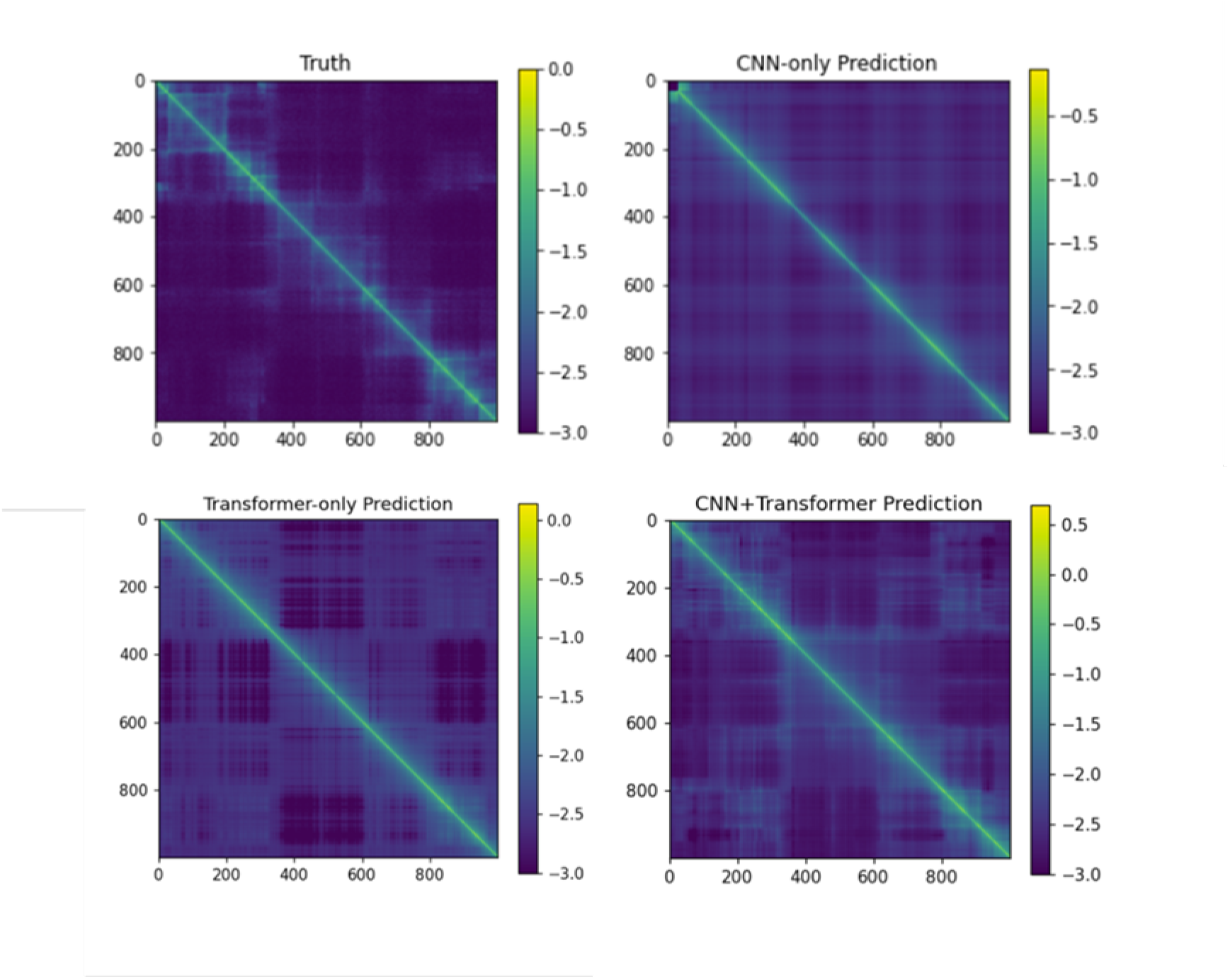
Optimization of the STS component. With curve-based normalization, STS with 8 CNN layers, STS with 8 transformer layers, and STS with 4 CNN layers and 4 transformer layers were added to UpDimConv STM, respectively. Their predicted matrices were compared to the true Hi-C matrix. The loss function was configured as *α*_1_ = 0.009, *α*_2_ = 0.001, *α*_3_ = 0.99, and all matrices were plotted on the log2 scale with pseudocount=0.001.

To ensure both stability and predicting accuracy, these models were saved when 80 training epochs were just finished, as too long training might result in model collapse with all outputs to be zeros.

### 3.4 Model Performance Analysis

Our final model was obtained after optimizing the STS composition and the loss formulation. With Pearsonr as the criteria to measure predicting accuracies, it outperformed previous methods, such as Akita (Pearsonr = 0.61) [2], DeepC (Pearsonr < 0.75) [3], HiC-Reg (Pearsonr around 0.6) [1], and Epiphany (Pearsonr = 0.7833) [4]. To explore the model generalization capability, test dataset of another cell line, GM12878, was generated using the same chromosomes as in the train dataset and test dataset of K562. Due to the high correlation of contact frequencies between these two cell lines (Figure 4a), the weight function (Method), which was previously used in K562 cell line, was also applied to GM12878. As a result, the model performed well on chromosome 18-22 and X with visible structural information (Pearsonr = 0.9226, Figure 4b), while the accuracy was slightly lower on chromosome 1-17 and the predicted matrices blurred more (Pearsonr = 0.9089, Figure 4c), which might be due to more complex interaction patterns on these chromosomes. Although the overall accuracies remained high on GM12878, it is still recommended to use HiCArch on the same cell line it has been trained. Alternatively, the weight function could be replaced by that of GM12878, but it required prior knowledge of Hi-C contact distribution of GM12878.

**Figure 4:**
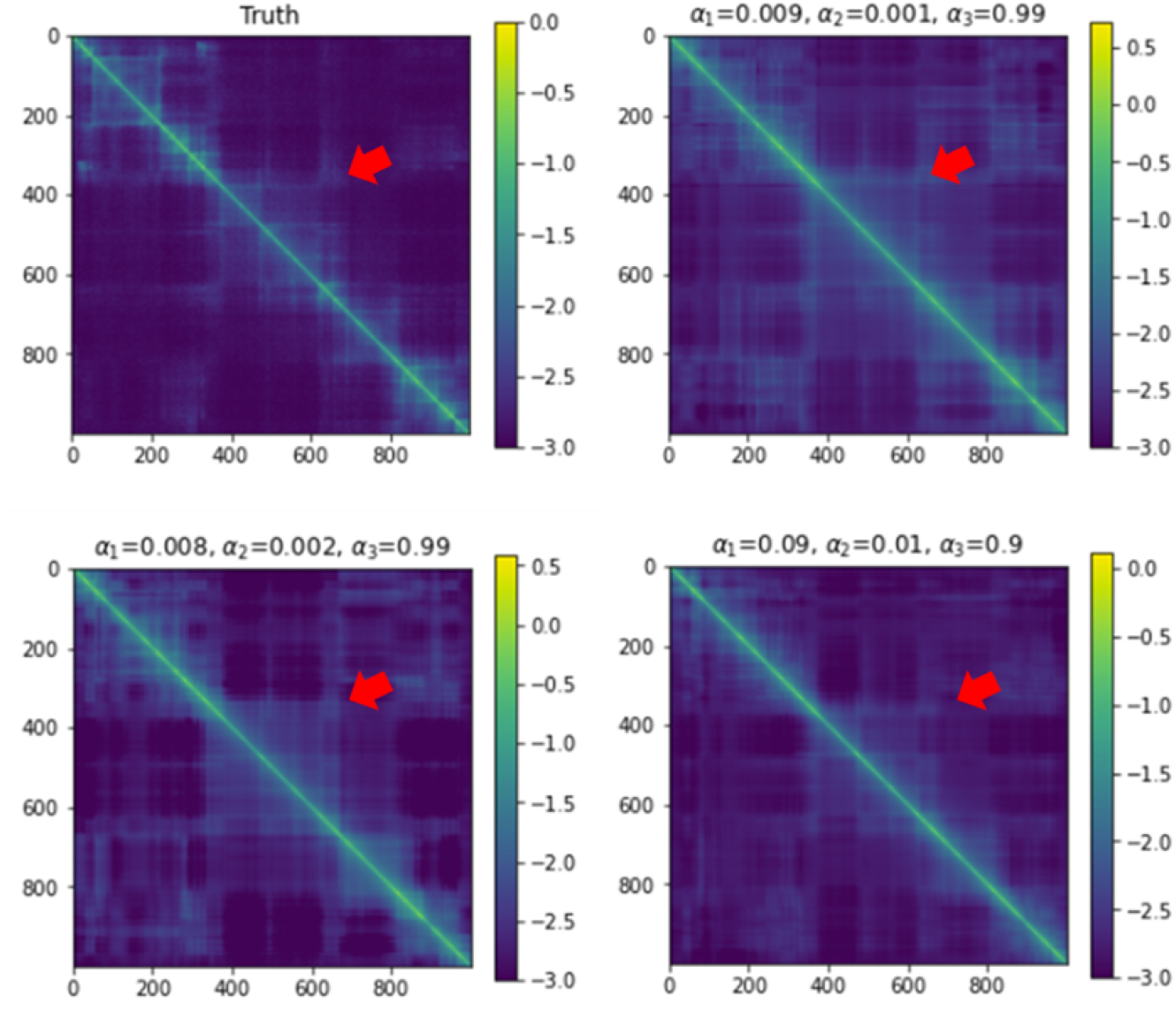
Calibration of term weights in loss formulation. The model used consisted of STS with 4 CNN layers and 4 transformer layers, and UpDimConv STM, as selected in the previous section. The matrices produced with varied *α*_1_, *α*_2_, *α*_3_ were compared to the true Hi-C matrix Red arrows indicate the position of a distal interaction in the true Hi-C matrix. All matrices were plotted on the log2 scale with pseudocount=0.001.

### 3.5 Epigenetic Feature Reduction

Until now, 11 epigenomic features were used as inputs in our model. However, it is also interesting to know which features are important than others so that the user could train their own model using the least epigenomic tracks but still with high predicting accuracies. Hence, HiCArch was trained on 10 features of K562 for 11 times, with deleting one epigenomic feature each time. While deleting other tracks made no differences, absence of H3K27ac, H3K4me3, H3K27me3, H3K9me3, H3K79me2 and TBP greatly reduced the test accuracies (Figure 5). Particularly, if the repressive marks H3K9me3 or H3K27me3 was unused, the model produced completely blank matrices even before being trained for 20 epochs. This result was partly consistent with HiC-Reg[1], which also recognized H3K27ac, H3K27me3, H3K9me3 as critical epigenetic marks. However, unlike HiC-Reg, H4K20me1, DNase I, H3K4me1 and CTCF were trivial in our experiments.

**Figure 5:**
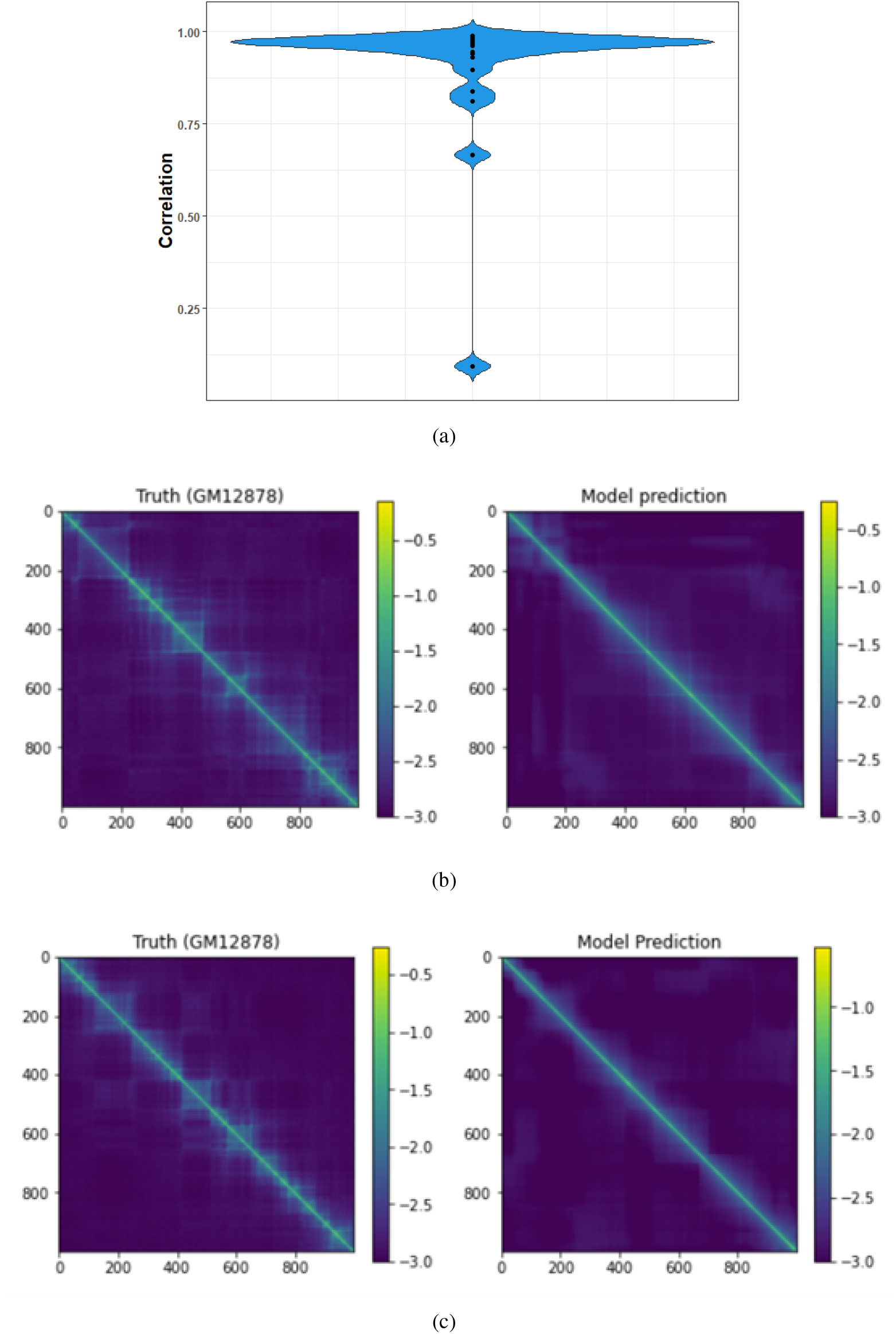
Generalization study on GM12878 cell line. Position-wise Pearson correlation between K562 and GM12878 on each chromosome was represented in the violin plot (a). The final model of HiCArch was tested on chromosome 18 to 22 and X (b) as well as chromosome 1 to 17 (c) of GM12878. All matrices were plotted on the log2 scale with pseudocount=0.001.

**Figure 6:**
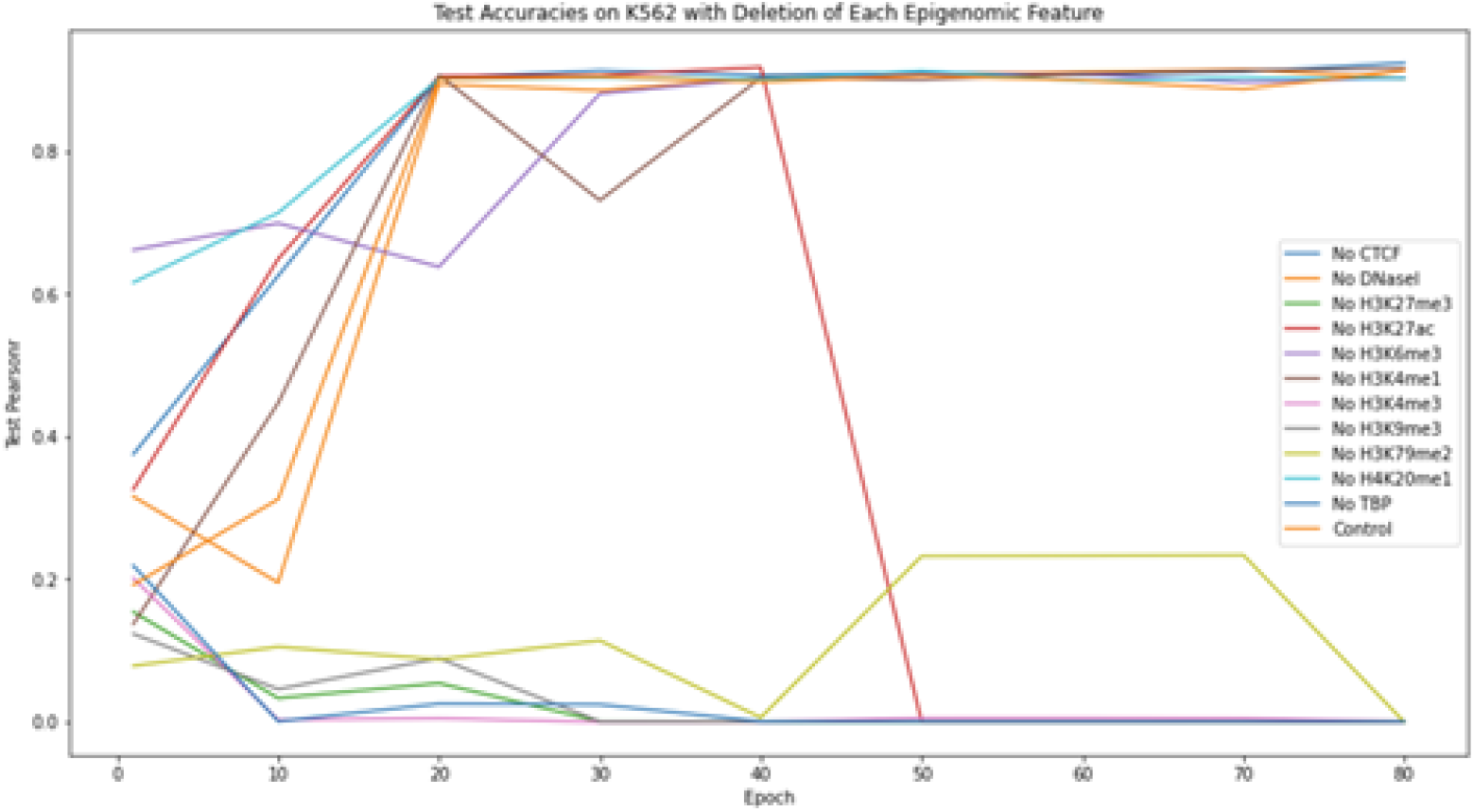
Feature reduction experiments with HiCArch. Each time one epigenomic feature was deleted, then the model was trained and tested on K562 with 10 remaining features. The curves represent the model test accuracies after being trained for 1-80 epochs.

## 4 Conclusion & Discussion

HiCArch is a light-weighted model that can accurately generate 2D Hi-C matrices from purely epigenetic tracks of the same resolution, without any input of exact DNA sequences. Also, HiCArch could recognize hierarchical genomics interactions, such as topological domains and distal contacts (up to 10Mb). It outperforms other existing tools on the Pearsonr accuracy metric, providing a more reliable approach to preview the genomic interactions before conducting actual Hi-C experiments. Although trained on K562, HiCArch could be generalized on other cell lines such as GM12878 with high accuracies, but the predicted matrices were not as clear as with original cell line. Therefore, the user could either train their own HiCArch with customized cell lines for more precise prediction, or try to substitute the weight function of K562 with the one generated with the target cell line. To further reduce users’ labor for data acquisition, model reduction experiments were done by identifying the important epigenetic features, and H3K27ac, H3K4me3, H3K27me3, H3K9me3, H3K79me2 and TBP were considered to have the greatest effect on HiCArch performance, which was partly consistent with previous research. HiCArch, even though harboring many advantages, is still unperfect now. For example, HiCArch requires at least 8Gb GPU memory for efficient training procedure, so we should further improve the model design in various ways, such as by trying more STS or STM components, so that HiCArch could be made more compatible to common PCs in the future.

## 5 Acknowledgement

We sincerely thank Prof. Chaochen Wang and Prof. Gaoang Wang of Zhejiang University International Campus for the hardware support of this work.

## Notes

https://www.encodeproject.org

### Competing Interest Statement

The authors have declared no competing interest.

